# Recruit symbiosis establishment and Symbiodiniceae composition influenced by adult corals and reef sediment

**DOI:** 10.1101/421339

**Authors:** A Ali, N Kriefall, LE Emery, CD Kenkel, MV Matz, SW Davies

**Affiliations:** Boston University, Biology Department, Boston, MA; The University of Texas at Austin, Department of Integrative Biology, Austin, TX; University of Southern California, Department of Biological Sciences, Los Angeles, CA

**Keywords:** coral, symbiosis, Symbiodiniceae, horizontal transmission, sediment, metabarcoding, ITS2

## Abstract

For most reef-building corals, the establishment of symbiosis occurs via horizontal transmission, where juvenile coral recruits acquire their algal symbionts (family Symbiodiniaceae) from their surrounding environment post-settlement. This transmission strategy allows corals to interact with a diverse array of symbionts, potentially facilitating adaptation to the newly settled environment. We exposed aposymbiotic *Pseudodiploria strigosa* recruits from the Flower Garden Banks to natal reef sediment (C-S+), symbiotic adult coral fragments (C+S-), sediment and coral fragments (C+S+), or seawater controls (C-S-) and quantified rates of symbiont uptake and Symbiodiniaceae community composition within each recruit using metabarcoding of the ITS2 locus. The most rapid uptake was observed in C+S+ treatments and this combination also led to the highest symbiont alpha diversity in recruits. While C-S+ treatments exhibited the next highest uptake rate, only one individual recruit successfully established symbiosis in the C+S-treatment, suggesting that sediment both serves as a direct symbiont source for coral recruits and promotes (or, potentially, mediates) transmission from adult coral colonies. In turn, presence of adult corals facilitated uptake from the sediment, perhaps via chemical signaling. Taken together, our results reinforce the key role of sediment in algal symbiont uptake by *P. strigosa* recruits and suggest that sediment plays a necessary, but perhaps not sufficient, role in the life cycle of the algal Symbiodinaceae symbionts.

## INTRODUCTION

Algal symbionts in the family Symbiodiniaceae are one of the most diverse groups of endosymbionts across marine environments and are hosted by a variety of invertebrates ranging from cnidarians, to mollusks, to sponges (Baker 2003; Stat et al. 2006; LaJeunesse et al. 2018). In adult tropical reef-building corals, these algal symbionts supply photosynthetic products to the coral host in return for inorganic nutrients and a residence (Muscatine and Porter 1977; Muscatine and Cernichiari 1969; Trench and Blank 1987). Coral-associated algal symbionts have radiated into genetically divergent lineages, formerly known as clades (A thru I: Stat et al., 2012) and recently reclassified as separate genera (LaJeunesse et al. 2018), which exhibit extensive morphological and functional diversity. Symbionts in the genus *Durusdinium* (clade D) have been shown to be highly infective (Abrego et al. 2009a) and offer increased thermal tolerance to their coral hosts (Berkelmans and van Oppen 2006) but reduce coral growth rates (Jones and Berkelmans 2011; Pettay et al. 2015). Alternatively, symbionts in the genus *Cladocopium* (clade C) provide a fitness advantage under ambient temperatures through increased carbon fixation and translocation (Cantin et al. 2009) and corals hosting algal symbiont in the genus *Symbiodinium* (clade A) experience reduced carbon fixation (Stat et al. 2008). Some corals have been shown to harbor a diverse assemblage of Symbiodiniaceae lineages, and it has been suggested that this functional diversity can greatly impact the ecology of a given coral host (*e.g.* Berkelmans and van Oppen 2006). It is also important to note that variation in thermal tolerance varies significantly within genera, such as *Cladocopium* (Howells et al. 2012) and interactions between hosts and symbionts also impact holobiont performance (Abrego et al. 2008; Cunning et al. 2015; Parkinson et al. 2015)

In corals, symbionts are either maternally transmitted (vertical transmission) or obtained from their environment (horizontal transmission) (Harrison and Wallace 1990; Baird et al. 2009). Corals that obtain their symbionts vertically are expected to host a lower diversity of symbionts since this relationship is stable through time, facilitating the co-evolution of host-symbiont partners (van Oppen 2004; Douglas 1998). On the other hand, horizontally transmitting species release aposymbiotic larvae that can travel great distances (Davies et al. 2015; Baums et al. 2014; Rippe et al. 2017) and upon settlement these recruits are capable of establishing symbiosis with diverse algal symbiont communities that do not necessarily reflect the symbionts hosted by local conspecifics or adults of their same species (Coffroth et al. 2001; Weis et al. 2001; Little et al. 2004; Abrego et al. 2009b). However, as coral recruits mature, the hosted symbiont community becomes dominated by a single clone of the lineage typical for the location (reviewed in Thornhill et al. 2017)) while establishment of symbiosis with novel Symbiodiniaceae species happens very rarely or never (Coffroth et al. 2010; LaJeunesse et al. 2010; Boulotte et al. 2016). Therefore, this initial acquisition of symbionts during recruitment represents a critical stage in coral-algal symbioses for horizontally-transmitting coral hosts.

The flexible symbioses of broadly dispersing, horizontally transmitting coral juveniles have been hypothesized to facilitate adaptation of the coral to environmental variation (Fournier 2013; van Oppen 2004; Sampayo et al. 2008; Davies et al. biorxiv), and indeed these associations have been implicated in local adaptation of the holobiont (Howells et al. 2013; Barfield et al. 2018). While much research has quantitatively described the diversity of coral-Symbiodiniaceae symbioses across species and environments at the adult life stage, much less is known about adaptations and mechanisms that symbionts employ to ensure transmission to the next coral generation. One potential mechanism for establishing symbiosis is through infection from a nearby conspecific adult coral (van Oppen 2004). Corals constantly expel photosynthetically active algal symbionts (Ralph et al. 2001; Hill and Ralph 2007). In theory, these cells could directly establish symbiosis with newly settled recruits. Alternatively, these expelled symbionts could colonize reef sediment, which could enable them to persist until the arrival of new recruits. Multiple studies have demonstrated that coral recruits are capable of establishing symbiosis in the presence of reef sediment (Adams et al. 2009; Cumbo et al. 2013; Nitschke et al. 2016), however it remains unclear whether these sediment-derived symbionts recapitulate the diversity of Symbiodiniaceae hosted by local adult corals on the same reef.

In this study, we first compared post-settlement symbiont uptake rates in the horizontally transmitting coral, *Pseudodiploria strigosa*, across multiple symbiont sources. *P*. *strigosa* recruits were placed in fully-crossed treatments that included the presence of natal reef adult coral fragments (C+S-), natal reef sediment (C-S+), a combination of adult coral fragments and natal reef sediment (C+S+), and seawater controls (C-S-) to test which environment promoted the most efficient uptake. The diversity of these established symbiont assemblages was examined using metabarcoding of the Internal Transcribed Region 2 (ITS2), to characterize Symbiodiniaceae communities within each individual recruit, adult coral fragment, and population of conspecific adults on the native reef to explore how variation in symbiont communities among recruits correlates with the Symbiodiniaceae communities found within local coral hosts.

## MATERIALS AND METHODS

### Experimental Methods

#### Coral spawning and larval rearing

During the annual coral spawning event at the Flower Garden Banks (FGB) on the evening of August 9^th^, 2012 at 21:15CDT (nine days after the full moon), gamete bundles from eight *Psuedodiploria strigosa* colonies were collected via scuba diving and spawning tents (Sharp et al. 2010). Gamete bundles were combined at the surface in a 14 L plastic tub filled with 1 μm filtered seawater (FSW) and left to cross-fertilize for two hours. Excess sperm was then removed by rinsing embryos through 150 μm nylon mesh. Developing larvae were reared in 1 μm FSW in three replicate plastic culture vessels at a density of two larvae per ml. Larvae were transferred to the laboratory at the University of Texas at Austin one-day post fertilization (dpf). Sediment collections were completed August 8^th^and were maintained in 1 μm filtered seawater. One large fragment of a single adult *Orbicella faveolata* was collected and maintained in the laboratory to serve as the adult coral source of algal symbionts. All collections were completed under the Flower Garden Banks National Marine Sanctuary (FGBNMS) permit #FGBNMS-2012-002.

#### Symbiont uptake experimental design

On August 14^th^, 2012 (5 dpf) twelve (5.5 gallon) experimental tanks were filled with artificial seawater (Instant Ocean, Blacksburg, VA, USA) and 800 ml of 1 μm filtered FGB water was added to each tank. Tanks were randomly assigned to one of four treatments (*n*=3/treatment): 1. FGB natal reef sediment only (C-S+), 2. *Orbicella faveolata* coral host fragment only (C+S-), 3. FGB natal reef sediment and *O. faveolata* coral host fragment (C+S+), and 4. Seawater control (C-S-) (Supplemental Figure S1). Tanks were maintained at identical salinity (35.5 ppt) and temperature (as measured by hobo data loggers: 25.5-28.5°C, Supplemental Fig. S2) throughout the uptake experiment. Four dpf, thousands of competent *P. strigosa* larvae were placed in sterile plastic dishes filled with artificial seawater (Instant Ocean, Blacksburg, VA, USA) and conditioned glass slides. Autoclaved, finely ground FGB crustose coralline algae (CCA) was added to slides to induce settlement (as per Davies et al. (2014, 2015)) and larvae were given four days in dark conditions to metamorphose. Four days later (8 dpf) plastic dishes were cleaned, and settlement conditions were replicated with new larvae to maximize recruitment rates per slide.

On August 21^st^ (12 dpf), slides with settled *P. strigosa* recruits were randomly placed into each treatment tank (*n*=3 slides per tank; Supplemental Figure S1). Symbiont uptake was visually assessed using a fluorescent stereomicroscope MZ-FL-III (Leica, Bannockburn, IL, USA) equipped with F/R double-bandpass filter (Chroma no. 51004v2). Recruits were considered as having established symbiosis when individual algal symbiont cells were obvious in recruit tentacles (Fig. 1 A, B). Recruits were surveyed daily from August 22-28^th^ (13-19 dpf), after which surveys were completed every three days. Uptake was continually monitored until October 16th (68 dpf) when final counts were completed due to algae overgrowth causing coral recruit death. Individuals successfully infected with symbionts were then individually collected using sterile razor blades, preserved in 95% EtOH and stored at -20°C until processing.

**Figure 1:**
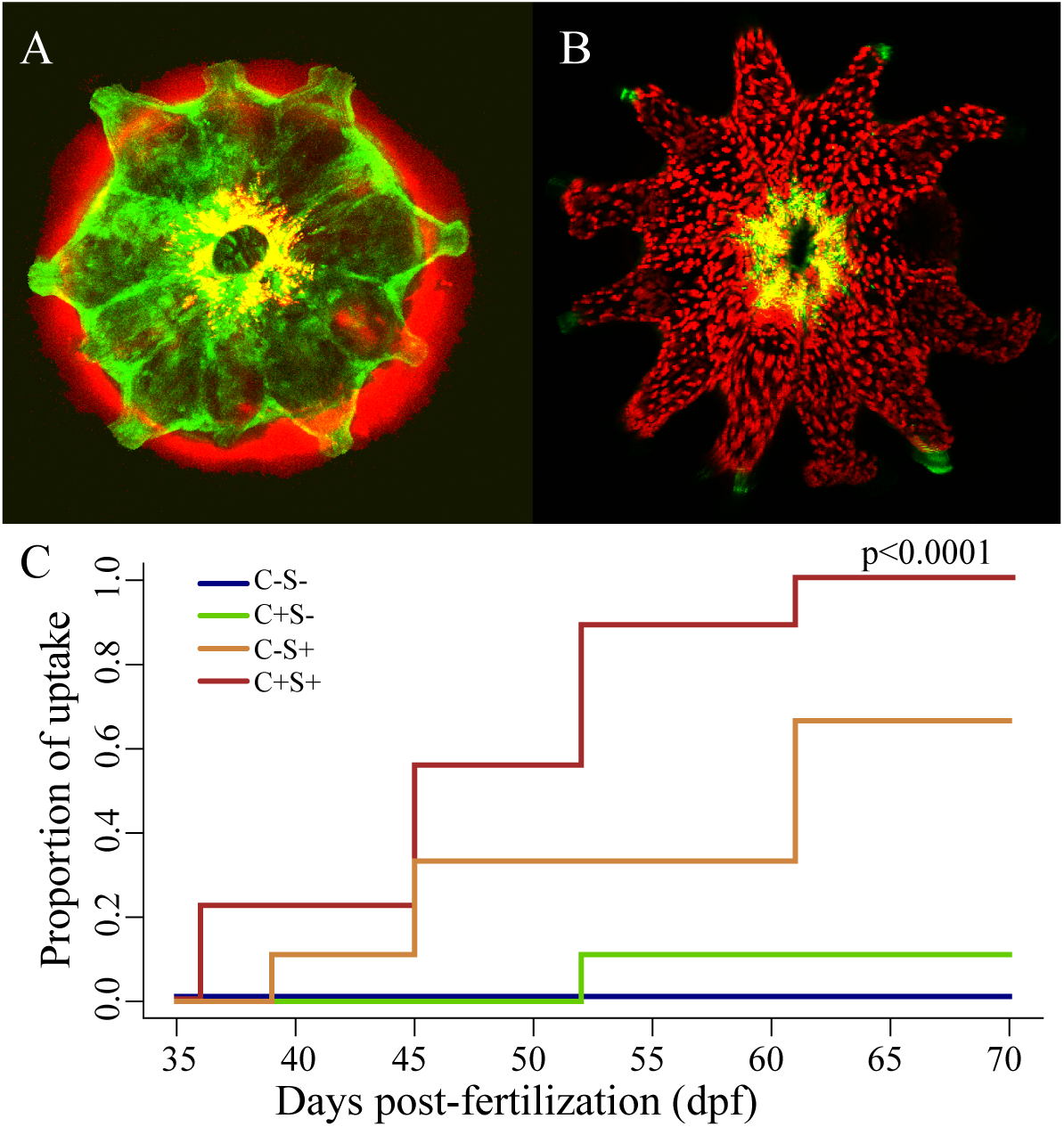
Algal symbiont uptake in *Pseudodiploria strigosa* recruits. Single *P. strigosa* recruit under confocal microscopy showing A. no algal symbiont uptake and B. symbiosis with the algal symbiont demonstrating the clear phenotypic differences in recruit uptake using fluorescence microscopy. Green fluorescence is innate green fluorescence from coral recruit and red excites chlorophyll, which can be seen surrounding the recruit in A (turf algae) and as discrete algal cells in B. C. Mean cumulative uptake of algal symbionts in *P. strigosa* recruits through time demonstrating the proportion of recruits that established symbiosis through time (dpf: days post fertilization) across the four experimental uptake treatments. P-value corresponds to cox-proportional hazards model indicating significant differences in uptake rate. C-S+ = FGB natal reef sediment only, C+S- = *Orbicella faveolata* coral host fragment only, C+S+ = FGB natal reef sediment and *O. faveolata* coral host fragment, and C-S- = seawater control.

#### Symbiont genotyping

Symbiont DNA was isolated from individual recruits using a DNeasy Plant Mini Kit (Qiagen) according to the manufacturer,s instructions. Recruits were disrupted by micropestle for 5 min using an aliquot of Lysing Matrix A (MP Biomedicals). Symbiont DNA was isolated from adult corals following Davies et al. (2013).

The ITS2 region was amplified via PCR using the forward primer *its-dino* (5, GTGAATTGCAGAACTCCGTG 3,) and the reverse primer *its2rev2* (5, CCTCCGCTTACTTATATGCTT 3,) (Pochon et al. 2001), following the protocols described in Kenkel et al. (2013) and Quigley et al. (2014), using 2μl of template DNA of unknown concentration. Briefly, amplifications were verified on agarose gels following 21 cycles and additional cycles were added as necessary to achieve a faint band (to reduce PCR biases) when 3μl of product was loaded on a 1% agarose gel and run for 15 min at 180 V (Supplemental Figure 3A). Cycle numbers ranged from 26 - 41 across samples (Supplemental Table S1), however several samples were amplified to 42 cycles along with no-template negative controls to assure that results at high cycle numbers were not due to contamination (Supplemental Figure 3B). These ,cycle-check, PCR,s were performed on a Tetrad 2 Peltier Thermal Cycler (Bio-Rad) using the following conditions: 94°C for 5 min, followed by 21 cycles of 94°C for 15 s, 59°C for 30 s and 72°C for 30 s and a final extension of 10 min at 72°C. Once optimal cycle numbers were obtained, all samples were re-amplified to their previously specified cycle number and verified on a gel to test for equivalent band intensity across samples (Supplemental Figure 3A).

Each PCR reaction was cleaned using a PCR clean-up kit (Fermentas) following the manufacturer,s instructions, measured using a NanoDrop 1000 spectrophotometer (Thermo Scientific) and diluted to 10 ng/μl. This product was then used as template for an additional PCR step used to incorporate 454-RAPID primers and barcodes to each sample. Each PCR contained 0.33 μM B-Rapid ITS2-forward primer (Br-ITS2-F: 5, CCTATCCCCTGTGTGCCTTGAGAGACGHC+GTGAATTGCAGAACTCCGTG 3,) in addition to 0.33 μM of unique A-Rapid-reverse primer containing an 8-bp barcode for subsequent sample identification (e.g. Ar-ITS2-R-16: 5, CCATCTCATCCCTGCGTGTCTCCGACGACT+**TGTAGCGC**+CCTCCGCTTACTTATATGCTT 3,, barcode sequence in bold). Each sample was uniquely barcoded.

Amplifications were visualized on a gel and based on visually assessed band intensities varying amounts of each barcoded sample were pooled for 454 sequencing. This pooled sample was cleaned via ethanol precipitation and re-suspended in 25 μl milli-Q water. 10 μl of this cleaned product was run on a 1% Agarose gel stained with SYBR Green (Invitrogen) for 45 min at 100V. The gel was visualized on a blue-light box and the target band was excised using a sterile razor blade and placed in 25 μl milli-Q water for overnight incubation at 4°C. The resulting supernatant was then submitted for 454 sequencing at the Genome Sequencing and Analysis Facility at the University of Texas at Austin. Raw sff files were uploaded to Sequence Read Archive (SRA) Accession Number SRP144167.

#### Statistical analyses

All analyses were completed in the R statistical environment (R Core Team 2017) and scripts are available at http://github.com/NicolaKriefall/sym_uptake. Rates of symbiont uptake by coral recruits were compared using the package *Survival* (Therneau and Lumley 2015). Numbers of recruits that established symbiosis with algal symbionts, measured as binary variables of successes and failures, were fit to a Cox,s proportional hazards regression model. A cumulative incidence curve was generated from this model and an ANOVA test was run to test for significant differences in uptake rates. To assess differences between pairs of treatments, the analysis was run pairwise for adult host fragment, natal reef sediment, and adult host fragment and natal reef sediment treatments.

To determine the community composition of Symbiodiniaceae in each coral recruit, 454 sequencing data were analyzed using the package *dada2* (Callahan et al. 2016). First, 454 pyrosequencing files were converted to FASTQ format using the package *R453Plus1Toolbox* (Klein et al. 2011) as *dada2* only processes FASTQ files (Callahan et al. 2016). Using *dada2*, FASTQ files were then trimmed to 300 bp in length as determined by associated quality profiles. Primers were clipped and sequences were de-replicated to obtain unique sequences. A sequence table was created to determine the distribution of sequence lengths in each sample and to remove those sequences that deviated from the expected sequence length. After de-noising sequencing data, chimeric sequences were removed and taxonomy was assigned by mapping to the GeoSymbio ITS2 database (Franklin et al. 2012). The package *phyloseq* (McMurdie and Holmes 2013) was then used to generate an OTU counts table (Supplemental Table S1) and to create bar plots to visualize and sort relative abundances of different Symbiodiniaceae lineages. Cumulative reads across lineages within a sample were then log-normalized following Green et al. (2014) and *pheatmap* (Kolde 2015) was used to visualize lineage differences across recruits and adults. *Phyloseq* was also used to construct an alpha diversity plot using Simpson and Shannon diversity controlling for effect of sample size. All raw sequence numbers through *dada2* filtering steps can be found in (Supplemental Table 2).

Lastly, a phylogenetic tree of the most abundant unique sequences from ITS2 sequencing of all samples (GenBank Accession #SUB4526136) together with a reference sequence of Symbiodiniaceae type B1 obtained from Green et al. (2014) and a second B1 reference and all other Symbiodiniaceae types from the GeoSymBio database by Franklin et al. (2012) was constructed. First, Multiple Sequence Comparison by Log-Expectation aligned sequences, Gblocks selected conserved sequences, phyML and the approximate Likelihood-Ratio Test (aLRT) assigned phylogeny and bootstrap values based on the maximum likelihoods model, and finally the TreeDyn function in Phylogeny.fr visualized the tree (Dereeper et al. 2008, 2010).The Newick output of the constructed tree was visualized using R package *ggtree* (Yu et al. 2017).

## RESULTS

### Coral recruit symbiont uptake

Initial symboint uptake by *P. strigosa* recruits was not observed until 36 days post fertilization (dpf), which was 18 days after recruits were added to uptake treatments. Uptake was confirmed by assessing chlorophyll fluorescence (Fig. 1A,B). The first recruits to exhibit uptake were in FGB natal reef sediment and *O. faveolata* coral host fragment treatments (C+S+) (Fig. 1C). Additionally, slides in C+S+ treatments were the only slides on which 100% of recruits successfully established symbiosis by the end of the experiment (68 dpf; 56 days after being placed in uptake treatments). The FGB natal reef sediment (C-S+) treatment was the second to exhibit uptake (Fig. 1C), however, significantly fewer recruits acquired symbionts when compared to C+S+ treatments (Wald,s *p* < 0.05). *Orbicella faveolata* coral host fragment (C+S-) treatments exhibited the slowest uptake rates (Fig. 1C), and final uptake proportions in this treatment were significantly lower than C+S+ treatments (Wald,s *p* < 0.01) and C-S+ treatments (Wald,s *p* < 0.05). As expected, recruits in seawater control treatments (C-S-) exhibited no uptake (Fig. 1C).

The likelihood of symbiont uptake by *P. strigosa* recruits was significantly affected by experimental treatment (*X*^2^= 30.779 and *p* < 0.0001). Hazard ratios from the Cox,s proportional hazards model demonstrated that uptake in the C-S+ treatment was significantly lower than the C+S+ treatment (0.294, CI: 0.097, 0.893), but higher than C+S-treatments (0.034, CI: 0.004, 0.283), suggesting that the presence of sediment increased the probability of symbiont acquisition in *P. strigosa* recruits.

### Symbiodiniaceae Genetic Diversity

To compare Symbiodiniaceae diversity among individual infected recruits across treatments, a total of 42 corals were successfully genotyped using 454 metabarcoding of the ITS2 locus (Supplemental Table S2). Thirty of these samples were individual recruits from experimental treatment tanks, six were *O. faveolata* host fragments from experimental treatment tanks and six were native *P. strigosa* adults collected from the east Flower Garden Banks (FGB) (Supplemental Table S1). A total of 67,027 raw reads were generated, 55,589 of which were left after adaptor trimming, quality filtering, and discarding reads shorter than 300 bp using the statistical package *dada2*. Recruit 2B1 was excluded from statistical analysis due to low number of remaining reads. Number of filter-passing reads in retained samples ranged from 589 to 4,341 with an average of 1264 reads (Supplemental Table S2).

The dominant lineage in adult *P. strigosa* was Symbiodiniaceae was B1 (genus *Breviolum* (LaJeunesse et al. 2018)), representing nearly 100% of sequences retrieved from *P. strigosa* colonies (*i.e.* they exclusively hosted B1, Fig. 2). Lineage B1 was also the dominant Symbiodiniaceae reference sequence in *O. faveolata* adults (98.7%-100%), but this species also associated with background levels of B10 (up to 1.3%, Fig. 2).

**Figure 2:**
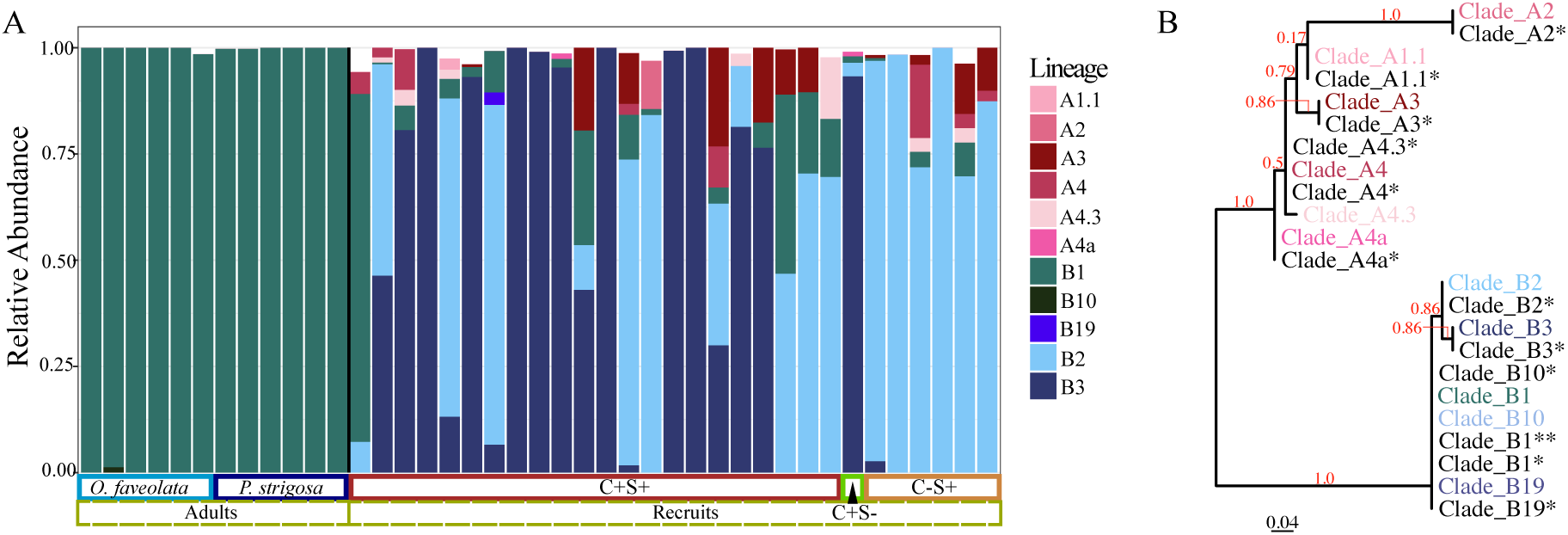
Symbiodiniaceae communities across adult *P. strigosa* in natal reef sites, adult *O. faveolata* used in uptake experiment (collected from natal reef site) and *P. strigosa* recruits in uptake experiment. A. Relative abundance of total reads mapping to reference sequences in GeoSymBio ITS2 database where each vertical bar denotes one coral,s Symbiodiniaceae community. The experimental uptake treatment of coral recruits and the two species of adult corals are indicated below the barplot. The black line demarcates adult corals from recruits, contrasting the differences in Symbiodiniaceae communities across the two life stages. C-S+ = FGB natal reef sediment only, C+S- = *O. faveolata* coral host fragment only, and C+S+ = FGB natal reef sediment and *O. faveolata* coral host fragment. B. Phylogenetic analysis of the most abundant unique Symbiodiniaceae sequences within coral samples in the present study in addition to reference ITS2 sequences that were successfully mapped to. Branch support values are shown on the branches at divisions between distinct clades in red. The scale bar represents replacements per nucleotide site. * indicates reference sequences from the GeoSymBio ITS2 database while ** indicates B1 reference sequence from Green et al. (2014).

Notably, the average proportion of lineage B1 in juvenile *P. strigosa* recruits was 8.9% (i.e. were background) and only a single recruit from the C+S+ treatment was dominated by B1 (Fig 2A). In general, the majority of sequences observed in coral recruits were not detected at any level in adult fragments (Fig 2A). Still, the two most common symbiont lineages among recruits did belong to the genus *Breviolum*, however they were lineage B2 (average proportion of 36.6% in recruits) and lineage B3 (average proportion of 43.5% in recruits). *P. strigosa* recruits also established symbiosis with a wider diversity of symbionts compared to adult samples (lineages: A1.1, A2, A3, A4, A4a, A4.3, B1, B10, B2, B19, B3; Fig 2 and Fig 3A). Lineage B2 was observed at higher abundances in C-S+ treatments, comprising an average proportion of 87.8% in each individual recruit, while B3 was common in treatments that included adult coral fragments (C+S- and C+S+). B3 represented 93.3% of sequences in C+S-treatment, however, these values were derived from a single recruit. In C+S+ treatments, lineage B3 was present at higher average proportions (53.3%) when compared to B2 (24.1%) (Fig 2; Fig 3A).

**Figure 3:**
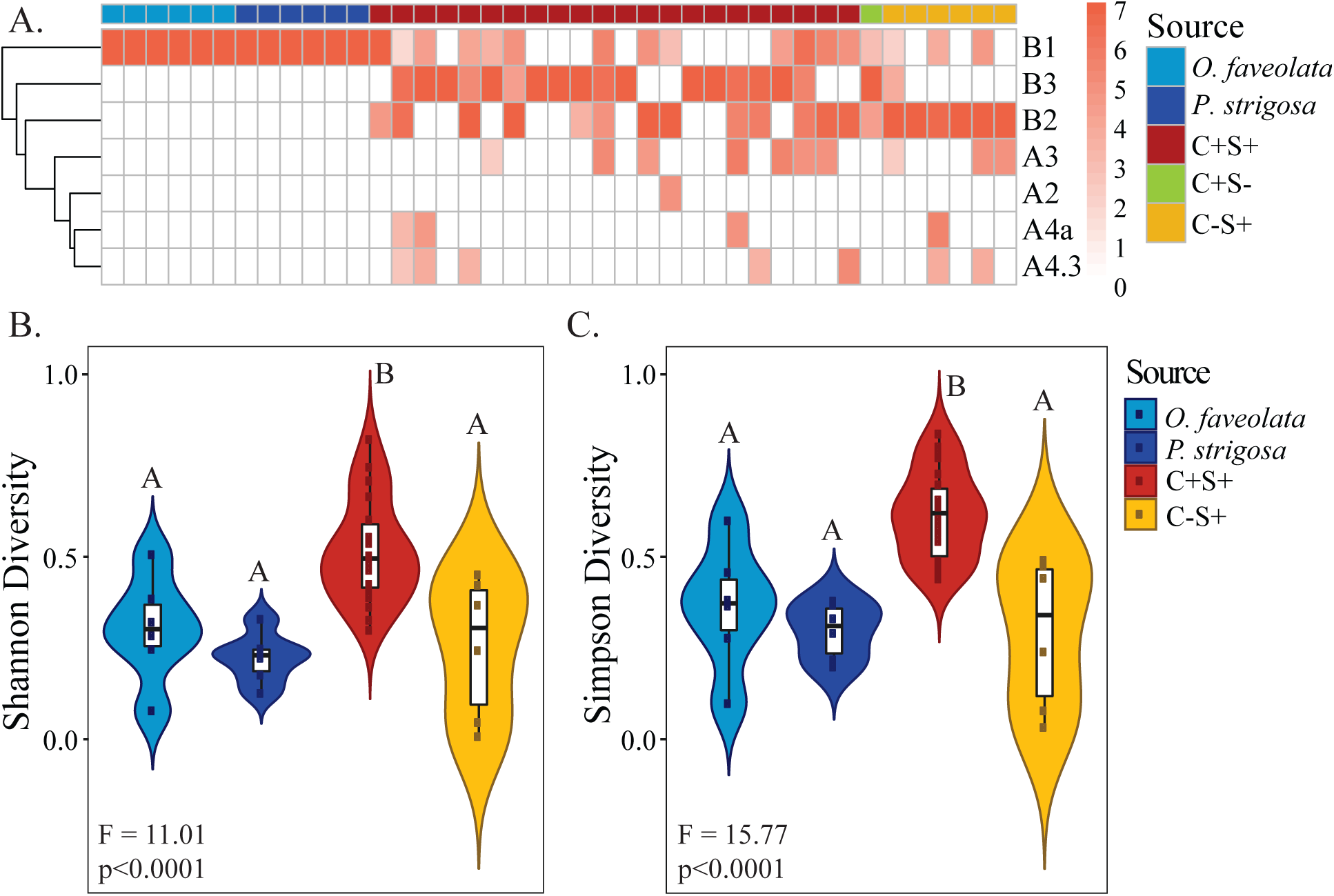
Symbiodiniaceae community diversity across adult *P. strigosa* in natal reef sites, adult *O. faveolata* used in uptake experiment (collected from natal reef site) and *P. strigosa* recruits in uptake experiment. A. Heatmap of *log* normalized counts within a lineage for all sequenced samples. B. Mean Shannon and Simpson alpha diversities for Symbiodiniaceae communities across adult corals and recruits in experimental uptake treatments. Widths of colored bands in violin plots correspond to the probability distribution of diversity indices. The boxplot and whiskers correspond to the interquartile range, median, and 95% confidence interval of alpha diversity measures. C-S+ = FGB natal reef sediment only, C+S- = *Orbicella faveolata* coral host fragment only, and C+S+ = FGB natal reef sediment and *O. faveolata* coral host fragment. *Orbicella faveolata* coral host fragment only treatments (C+S-) were excluded from analyses due to low sample size (N=1).

Shannon and Simpson alpha diversity were calculated for each treatment and a one-way ANOVA tested for diversity differences across symbiont source treatments. Both diversity measures indicated that alpha diversity varied significantly across source treatments (Simpson: F=15.77, *p*<0.0001; Shannon: F=11.01, *p*<0.0001; Table 1) and Tukey,s post-hoc tests confirmed that C+S+ treatments exhibited significantly higher mean Simpson and Shannon alpha diversities when compared to alpha diversities of other treatments and adult host fragments of both species (*p*<0.05; Table 2). C+S-treatment was not included in pairwise comparisons since only a single recruit achieved symbiosis.

**Table 1:** Tukey post-hoc pairwise statistics for Simpson and Shannon alpha diversities with respect to recruits in different uptake treatments and adult coral host Symbiodiniceae communities. P-values: *<0.05, **<0.01, ***<0.001 Note: Adult host fragment only treatments (C+S-) were not included given that too few recruits were observed to uptake algal symbionts. C-S+ = FGB natal reef sediment only and C+S+ = FGB natal reef sediment and *O. faveolata* coral host fragment.

| Treatment | Simpson p-value | Shannon p-value |
| --- | --- | --- |
| Adult *O. faveolata* – *P. strigosa* | 0.8404 | 0.7645 |
| Adult *P. strigosa* – C+S+ | 0.0017** | 0.0124 * |
| Adult *P. strigosa* – C-S+ | 0.8119 | 0.9402 |
| Adult *O. faveolata* – C+S+ | 0.0001*** | 0.0004 *** |
| Adult *O. faveolata* – C-S+ | 0.9999 | 0.9773 |
| C-S+ – C+S+ | 0.0001*** | 0.0018 ** |

## DISCUSSION

The reservoirs of free-living Symbiodiniaceae available for uptake by horizontally transmitting corals remain unresolved (Quigley et al. 2017). Here we assessed the relative roles that availability of reef sediment and coral adults play in the establishment of symbiosis in the horizontally transmitting reef-building coral *Pseudodiploria strigosa*. We found that reef sediment appears necessary for the successful establishment of symbiosis in *P. strigosa* coral recruits since recruits in treatments with sediment (C+S+ and C-S+) consistently exhibited significantly higher uptake rates when compared to treatments without sediment (C+S- and C-S-) (Fig 1C). This outcome is consistent with previous studies investigating symbiont uptake, which have similarly found that sediment serves as an important reservoir of Symbiodiniaceae for horizontally transmitting coral larvae and recruits (Adams et al. 2009; Cumbo et al. 2013; Nitschke et al. 2016). Free-living Symbiodinaceae are ubiquitous in the reef environment (Coffroth et al. 2006; Pochon et al. 2010; Takabayashi et al. 2012; Quigley et al. 2017; Porto et al. 2008) and their densities in the sediment have been estimated to be up to 15 times higher when compared to densities in the water column (Littman et al. 2008) due to the symbiont,s largely immobile lifestyle and because they are negatively buoyant (Coffroth et al. 2006; Yacobovitch et al. 2004). In light of these Symbiodiniaceae distributions, it is perhaps not surprising that we observed significantly higher uptake rates in treatments with sediment available (C-S+, C+S+) (Fig 1A).

We observed that only a single coral recruit took established symbiosis in the presence of adult coral but in the absence of sediment (C+S-); notably, most of these symbiont types were not detected in the adult tissue (Fig 2). Therefore, similarly to the algal communities established from sediment, these symbiont lineages likely represent free-living strains populating the coral,s surface or exposed skeleton rather than algal symbionts establishing symbiosis directly from adult coral.

The most surprising of our results is the finding that the combined C+S+ treatment exhibited much higher uptake rates than can be expected from just the sum of individual C-S+ and C+S-effects (Fig 1). Perhaps the presence of an adult coral alone increases uptake rates through the use of chemical cues. Previous work has linked chemical cues between corals and algal symbionts (Fitt and Trench 1981; Hagedorn et al. 2015; Fitt 1984; Takeuchi et al. 2017), however facilitation of the onset of symbiosis via adult specific cues is a novel hypothesis. In turn, the presence of sediment might facilitate uptake of algal symbionts from other sources. If the B3 symbiont type, which was never detected in adult coral colonies and was detected nearly exclusively in C+ treatments, is derived from the surface of the adult coral, then the sediment appears to have strongly promoted its uptake (Fig 2A). It is possible that sediment is required for the alga to complete a certain life cycle transition before it can infect recruits.

While *P. strigosa* recruits took up a small proportion of the “adult-like” B1 symbiont, they also took up many other Symbiodiniaceae lineages that were undetectable in adult *O. faveolata* (Fig 2A; Fig 3A). Stark differences in Symbiodiniaceae communities between early life stages and adults have been observed in multiple horizontal-transmitting corals, including Pacific Acroporids (Abrego et al. 2009a, 2009b; Little et al. 2004; Gómez-Cabrera et al. 2007) and Caribbean *Orbicella faveolata* (McIlroy and Coffroth 2017). We demonstrate similar results for a divergent Caribbean coral lineage (*P. strigosa*), suggesting that this phenomenon is a common feature of horizontally transmitting corals.

We did not assess the Symbiodiniaceae diversity present in the sediment, so we cannot determine if uptake of symbionts from the sediment was random or if certain lineages were more infectious. Future studies should sequence sediment Symbiodiniaceae communities to address this shortcoming, especially given that Quigley et al. (2017) found that Symbiodiniaceae communities in the sediment had four times as many OTUs when compared with Symbiodiniaceae communities hosted by juvenile *Acropora* recruits. In addition to finding more OTUs, Quigley et al. (2017) determined that very few OTUs were shared among juveniles and sediment, indicating that infection capabilities of different strains are not equal.

While high diversity symbiont communities in juvenile horizontally transmitting corals are well-established, the reason for this remains unclear. Increased diversity in recruits could be due to the lack of robust symbiont recognition mechanisms (Cumbo et al. 2013). Alternatively, harboring a more diverse Symbiodiniaceae community could confer varied functional and physiological advantages, perhaps even allowing them to cope with a variable local environments (Thornhill et al. 2017). Interestingly, uptake of B1 was not significantly higher in the presence of the coral fragment, suggesting that endosymbiotic B1 cells did not contribute significantly to the algal population capable of infecting recruits. Once again, this suggests that infective Symbiodiniaceae cells are free-living, and that transition to this state from the state of endosymbiosis is either indirect or takes considerable time.

Our results corroborate prior work showing that both sediment and host corals enhance the establishment of symbiosis in horizontally-transmitting corals. Most notably, we found that the presence of adult corals interacted synergistically with the presence of sediment. Clearly, more work on the life history of Symbiodiniaceae is required to explain these observations and to understand all the steps leading to transmission of resident endosymbionts to the next generation of coral hosts.

## Supplemental Information

**Supplemental Figure 1:**
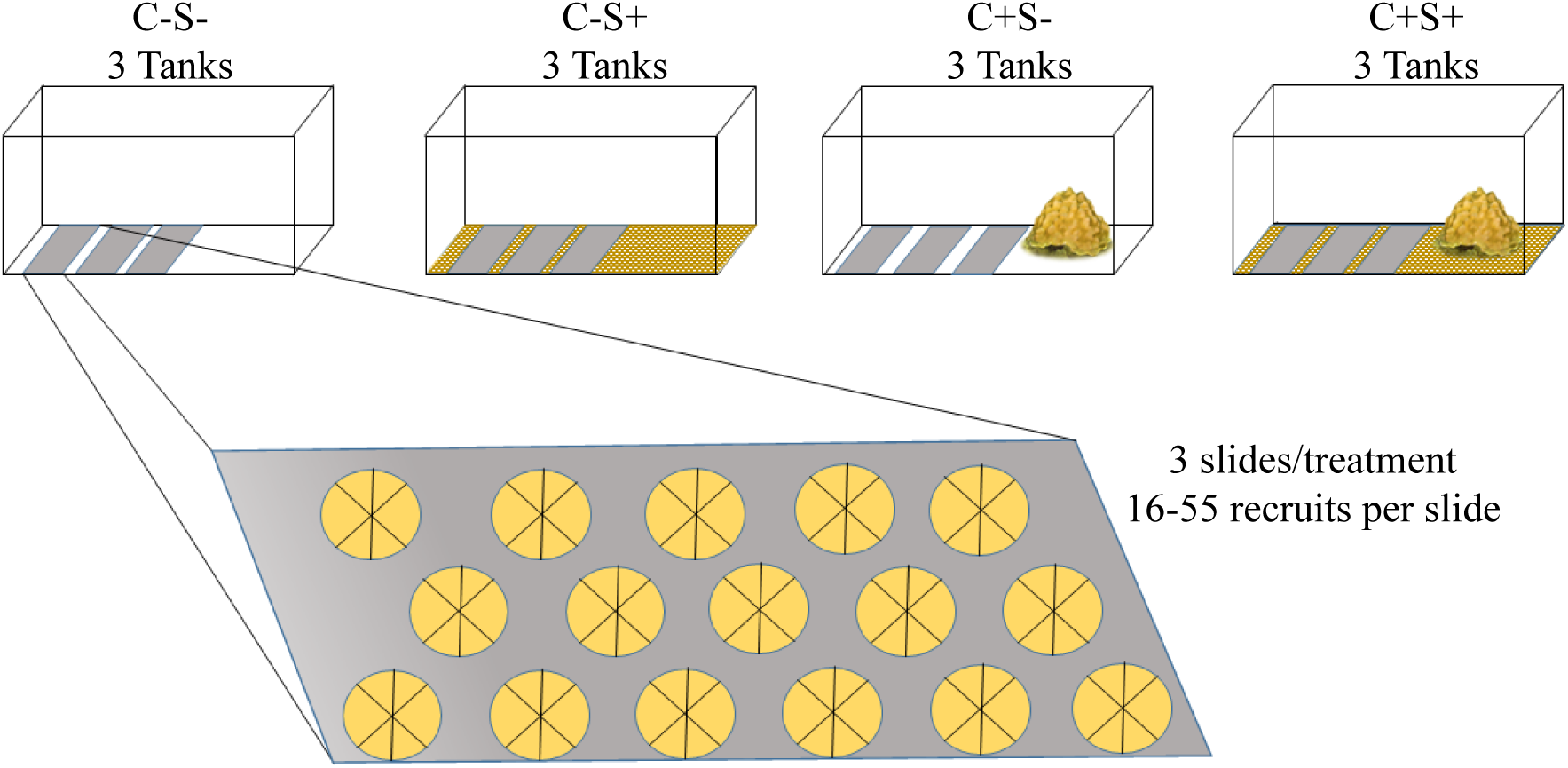
Symbiodiniaceae uptake experimental design demonstrating the four different uptake treatments and the three replicate slides with settled *P. strigosa* corals (N=16-55 recruits per slide) within each tank (3 tank systems for each uptake treatment). C-S+ = FGB natal reef sediment only, C+S- = *Orbicella faveolata* coral host fragment only, C+S+ = FGB natal reef sediment and *O. faveolata* coral host fragment, and C-S- = seawater control.

**Supplemental Figure 2:**
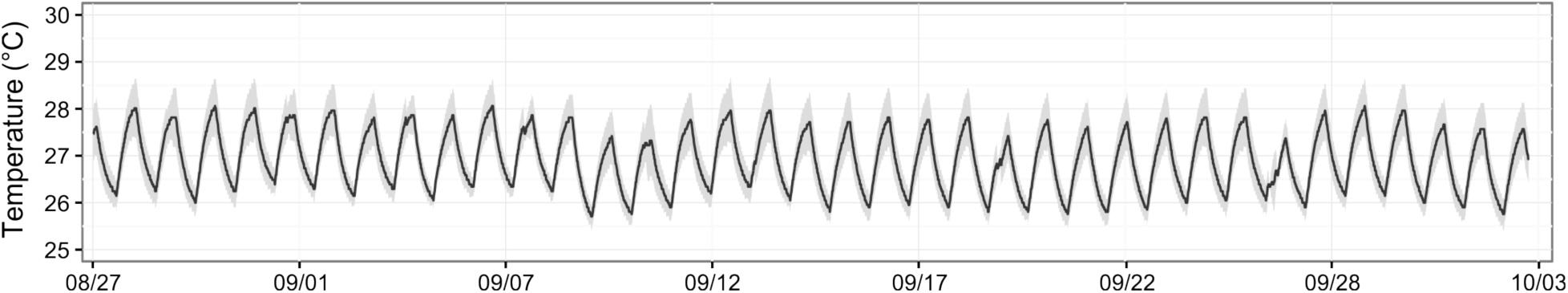
Mean temperature in a representative experimental tank through time. Black line indicates mean temperature and grey shading shows 95% confidence interval around that mean. Data were collected using Hobo data loggers.

**Supplemental Figure 3:**
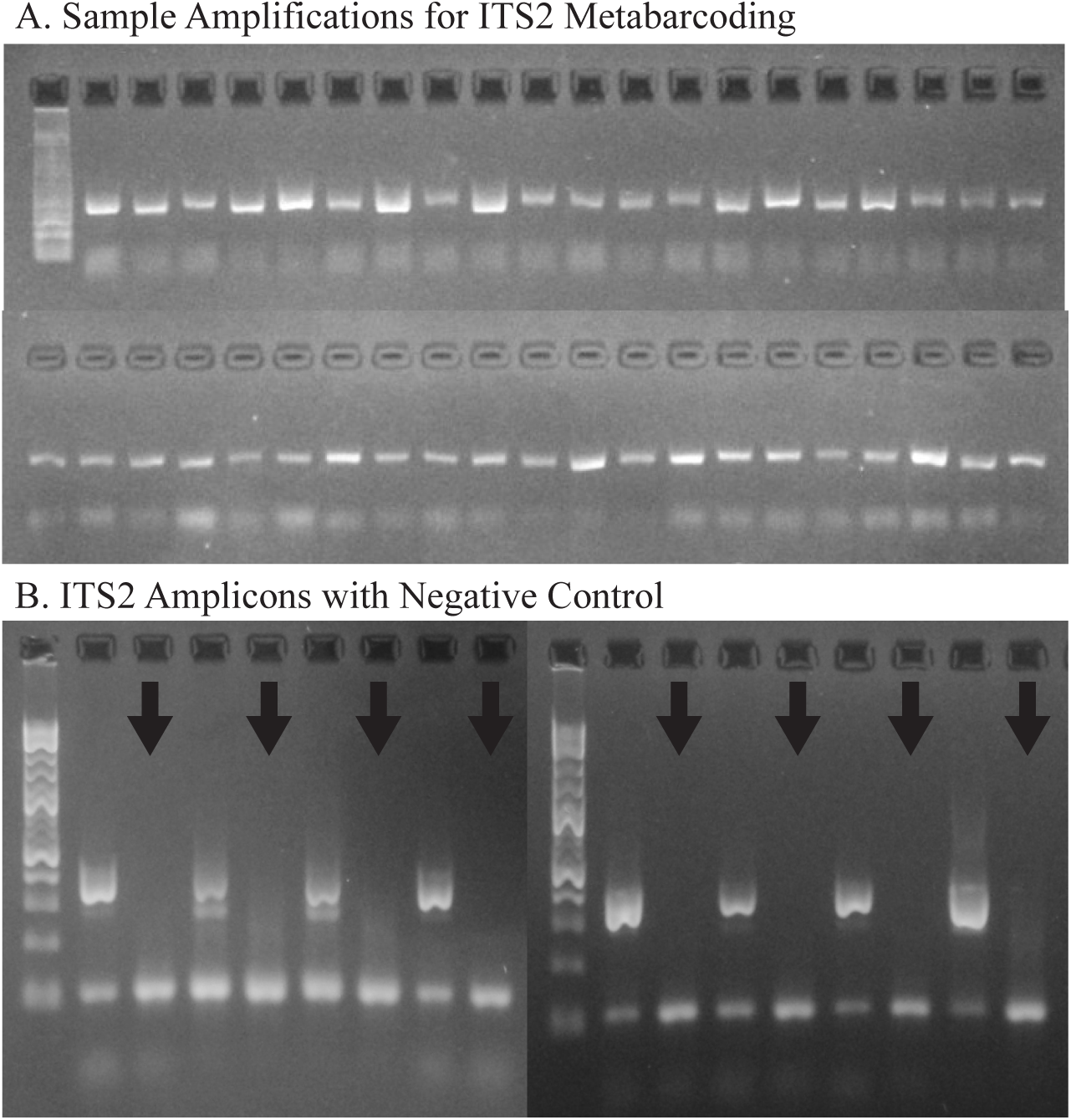
ITS2 Symbiodiniaceae community library preparation. A. ITS amplicon for each sequenced coral adult and *P. strigosa* recruit. Cycle numbers ranged from 26 - 41 across samples (Supplemental Table S1). B. ITS2 amplicons alongside their no-template negative controls (black arrows) demonstrating that even under high cycle numbers (42 cycles) no amplification is observed in negative controls.

Supplemental Table S1: Raw OTU counts

**Supplemental Table S2:**
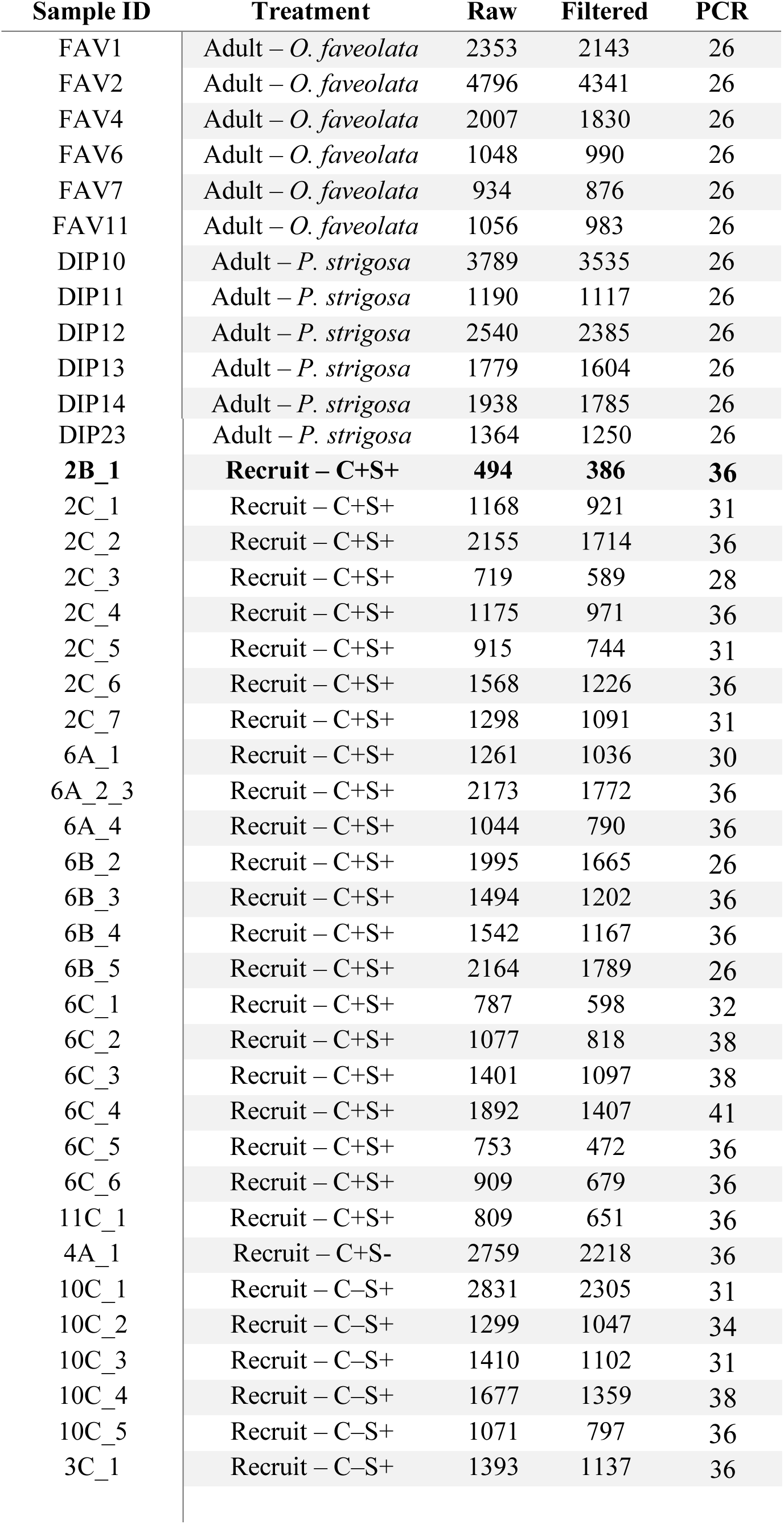
Coral adult and recruit DNA sample ID,s and their associated uptake treatment tanks, raw 454 sequence numbers, *dada2* filtered sequence numbers and total number of PCR cycles (PCR) ran for each sample to achieve a visual band on agarose gel (Supplementary Figure 3A). Sample in bold was not included in downstream analyses due to low read depth. C-S+ = FGB natal reef sediment only, C+S- = *Orbicella faveolata* coral host fragment only, C+S+ = FGB natal reef sediment and *O. faveolata* coral host fragment, and C-S- = seawater control.

